# Impact of sequencing depth and read length on single cell RNA sequencing data: lessons from T cells

**DOI:** 10.1101/134130

**Authors:** Simone Rizzetto, Auda A. Eltahla, Peijie Lin, Rowena Bull, Andrew R. Lloyd, Joshua W. K. Ho, Vanessa Venturi, Fabio Luciani

## Abstract

Single cell RNA sequencing (scRNA-seq) has shown great potential in measuring the gene expression profiles of heterogeneous cell populations. In immunology, scRNA-seq allowed the characterisation of transcript sequence diversity of functionally relevant sub-populations of T cells, and notably the identification of the full length T cell receptor (TCRαβ), which defines the specificity against cognate antigens. Several factors, such as RNA library capture, cell quality, and sequencing output have been suggested to affect the quality of scRNA-seq data, but these factors have not been systematically examined.

We studied the effect of read length and sequencing depth on the quality of gene expression profiles, cell type identification, and TCRαβ reconstruction, utilising 1,305 publically available scRNA-seq datasets, and simulation-based analyses. Gene expression was characterised by an increased number of unique genes identified with short read lengths (<50 bp), but these featured higher technical variability compared to profiles from longer reads. TCRαβ were detected in 1,027 cells (79%), with a success rate between 81% and 100% for datasets with at least 250,000 (PE) reads of length >50 bp.

Sufficient read length and sequencing depth can control technical noise to enable accurate identification of TCRαβ and gene expression profiles from scRNA-seq data of T cells.

## INTRODUCTION

Single cell RNA sequencing (scRNA-seq) has vastly improved our ability to determine gene expression and transcript isoform diversity at a genome-wide scale in different populations of cells. scRNA-seq is becoming a powerful technology for the analysis of heterogeneous immune cells subsets ^1,2^ and studying how cell-to-cell variations affect biological processes ^3,4^. Despite its potential, scRNA-seq data are often noisy, which are caused by a combination of experimental factors, such as the limited efficiency in RNA capture from single cells, and also by analytical factors, such as the challenges in separating true variation from technical noise ^5-7^. The quality of scRNA-seq data depends on mRNA capture efficiency ^8^, the protocol utilised to obtain libraries, as well as sequence coverage and length ^3,4^. Bioinformatics tools for the analyses of scRNA-seq data have been rapidly evolving, whereby various algorithms have been proposed to resolve the issues related to scRNA-seq compared to classical bulk transcriptomic analysis ^9,10^. However, the lack of a consensus in the data analyses further contributes to difficulties in assessing the quality of the data analysed so far.

One important consideration in designing scRNA-seq experiments is to decide on the desired sequencing depth (*i.e.*, the expected number of reads per cell) and read length ^3,6^. These are two important experimental parameters that can be controlled, and which need to be often predetermined before sequencing. For bulk RNA-seq data, sequencing depth and read length are known to affect the quality of the analysis ^11^. For scRNA-seq it has been shown that half a million reads per cell are sufficient to detect most of the genes expressed, and that one million reads are sufficient to estimate the mean and variance of gene expression ^6^. Low coverage scRNA-seq has also been utilised to show that 50,000 reads per cell are sufficient to classify a cell type in a sample of 301 cells ^12^. Nevertheless, this may not be sufficient when more homogenous populations are involved, for example T cell subsets, such as central memory and effector memory cells. In these scenarios, deep sequencing of single cell library may be required for improving detection of genes with low expression ^3,6^. Indeed, an important issue for scRNA-seq data is the very large number of genes with no detectable expression in a cell ^6^. This overrepresentation of zeros in scRNA-seq datasets makes it difficult to distinguish technical dropout of transcripts from true biological variation between cells ^3^.

Nonetheless, there has not been any systematic evaluation of the effect of sequencing depth and read length on scRNA-seq data analysis. In designing a scRNA-seq experiment it is optimal to generate data by maximising sequencing depth and utilising the longest read length. This approach would improve the quality of the reads alignment and also maximise the chance of detecting low abundant transcripts. In reality, we are often constrained by the cost of sequencing. Therefore a more practical question is to ask what is the minimum sequencing depth and read length that allows users to obtain adequate information for their desired downstream analyses.

To answer these questions, we have focussed on assessing the quality of available scRNA-seq data from T cells, which form a highly heterogeneous population of lymphocytes that play a vital role in mounting successful adaptive immune responses against intracellular pathogens and tumours ^13^. T cells are also characterised by a highly diverse repertoire of TCRs, which identify the specific recognition of the cognate antigen. TCRs are heterodimer proteins composed of two chains, α and β, and a subset of those expressing the γδ chains, which result from genetic recombination of the V(D)J genes. The diversity of TCRαβ repertoire has been associated with successful control of many pathogens ^14^, and more recently with outcome of checkpoint inhibitor immunotherapy for patients with metastatic melanoma ^15^. The highly polymorphic nature of the TCR genes has made their identification very difficult in bulk population sequencing datasets. In the last decade, deep sequencing approaches of bulk TCRs focusing on either α or β chains ^16^. The advent of scRNA-seq allowed the identification of the full length TCR of both α and β chains (referred to hereafter as TCRαβ) from T cells ^17,18^. This has now led to the capacity to simultaneously detect TCRαβ and full gene expression profiles in one experiment, thereby allowing direct study of TCR diversity and its interaction with the T cell functions reflected in gene expression profiles.

In this study we performed a comprehensive analysis of the impact of sequencing depth and read length on the detection of full length TCRαβ sequences, as well as estimation of gene expression and its effect on cell-type identification. Our study aims to fill this gap through performing a re-analysis of eight published scRNA-seq data that have a wide range of read length and sequencing depth, and analysis of simulated datasets that were subsampled from a deeply sequenced human T-cell scRNA-seq dataset. The analysis suggests important precautionary steps for researchers seeking to maximise throughput of single cell experiments without compromising the quality of the results.

## RESULTS

To assess the effects of sequencing depth and read length on accurate reconstruction of full length TCRαβ and gene expression profile from scRNA-seq data, we manually reviewed NCBI’s Gene Expression Omnibus ^19^ and ArrayExpress ^20^ to identify relevant T-cell scRNA-seq data published prior to April 2016. Eight datasets were identified with accessible data, collectively profiling 1,305 single cells (Table 1). The datasets were generated from mouse ^18,21-23^ and human-derived cells ^17^, utilising one of the available versions of the Smart-Seq protocol ^24^, and had a wide range of sequencing depth (1.2-8.4 million paired-end (PE) reads per cell) and read length (25-215bp) (Table 1). The number of expressed genes identified among the 1,305 cells ranged between 2,563 and 6,795 per cell (Table 1). We observed a weak negative linear correlation between read length and the average number of genes identified within a dataset (R=-0.49), but no relationship with the number of genes identified in each cell (R=-0.16). There was no clear correlation between the number of genes identified and the sequencing depth (R=0.1).

**Table 1.** scRNAseq data sets analysed in this study.

| **Dataset** | **Dataset** | **Ref** | **Accession number** | **Organism** | **Number of cells** | **Average reads length (nt)** | **Average number of PE reads (x10<sup>6</sup> reads)** | **scRNA-seq protocol** | **Number of genes expressed (FPKM>1) per cell** |
| --- | --- | --- | --- | --- | --- | --- | --- | --- | --- |
| 1 | HCV specific CD8+ T cells | <sup>33</sup> | E-MTAB-4850 | human | 54 | 145 | 8.4 | Smart-Seq2 | 2,563 |
| 2 | HCV specific CD8+ T cells | <sup>33</sup> | E-MTAB-4850 | human | 12 | 215 | 3.5 | Smart-Seq2 | 3,289 |
| 3 | Th17 cells (A) | <sup>21</sup> | GSE74833 | mouse | 399 | 125 | 2.5 | Smart-Seq | 5,128 |
| 4 | Th17 cells (B) | <sup>21</sup> | GSE74833 | mouse | 269 | 100 | 3.7 | Smart-Seq | 6,540 |
| 5 | Th17 cells (C) | <sup>21</sup> | GSE74833 | mouse | 100 | 25 | 1.5 | Smart-Seq | 4,146 |
| 6 | CD4+ T cells | <sup>18</sup> | E-MTAB-3857 | mouse | 272 | 100 | 4.3 | Smart-Seq | 2,354 |
| 7 | CD8+ T cells | <sup>34</sup> | GSE74923 | mouse | 106 | 32 | 1.2 | Smart-Seq2 | 6,796 |
| 8 | Th2 | <sup>23</sup> | E-MTAB-2512 | mouse | 93 | 75 | 1.6 | Smarter-Seq | 6,401 |

### The effect of sequencing depth and read length on reconstruction of full-length T-cell receptors

We analysed whether sequencing depth and read length affect the detection and reconstruction of TCRαβ. Two recently developed bioinformatics methods for reconstruction of full-length TCRαβ from scRNA-seq data were used, TraCeR ^18^ and VDJPuzzle ^17^. The analysis performed with VDJPuzzle and TraCeR revealed successful TCRαβ reconstruction in 1027 cells (79%) and 953 cells (73%), respectively (Table 2). With VDJPuzzle consistently had a higher reconstruction rate than TraCeR, with regard to reconstruction of the αβ full length, as well as of the single chain repertoires (*i.e.*, α or β) (*p*<0.05; paired t-test; Table 2). Six of the eight datasets had a success rate >80% in detection of TCRαβ, and up to 100% for scRNA-seq datasets with an average read length of 215 bp. The two datasets with lowest detection rate of TCRαβ had 25 and 32 bp long reads, where only 0% and 1.89% of the cells successfully generated TCRαβ sequences, respectively (Table 2 and Fig. 1A). In terms of sequencing depth, datasets with less than 0.25 million PE reads resulted in detection of TCRαβ in less than 1% of the cells, and this increased rapidly to >80% for depths >0.25 million PE reads (Fig. 1B).

**Table 2.** The success rate of reconstructing full-length T-cell receptors (TCR) using VDJPuzzle and TraCeR for the various scRNA-seq data sets.

| **Dataset** | **Number of cells** | **Average reads length (nt)** | **Average number of PE reads ($\times 10^6$ reads)** | **TCRα success rate (%) VDJPuzzle** | **TCRβ success rate (%) VDJPuzzle** | **TCR$\alpha\beta$ success rate (%) VDJPuzzle** | **TCRα success rate (%) TraCeR** | **TCRβ success rate (%) TraCeR** | **TCR$\alpha\beta$ success rate (%) TraCeR** |
| --- | --- | --- | --- | --- | --- | --- | --- | --- | --- |
| 1 | 54 | 145 | 8.4 | 81.48 | 85.19 | 81.48 | 81.48 | 85.19 | 81.48 |
| 2 | 12 | 215 | 3.5 | 100.00 | 100.00 | 100.00 | 100.00 | 100.00 | 100.00 |
| 3 | 399 | 125 | 2.5 | 99.25 | 98.75 | 98.50 | 92.98 | 93.48 | 91.23 |
| 4 | 269 | 100 | 3.7 | 98.51 | 98.88 | 97.77 | 89.59 | 94.42 | 86.62 |
| 5 | 100 | 25 | 1.5 | 0.00 | 0.00 | 0.00 | 0.00 | 0.00 | 0.00 |
| 6 | 272 | 100 | 4.3 | 89.71 | 93.38 | 85.66 | 80.88 | 91.54 | 80.88 |
| 7 | 106 | 32 | 1.2 | 1.89 | 7.55 | 1.89 | 2.83 | 0.00 | 0.00 |
| 8 | 93 | 75 | 1.6 | 89.25 | 93.55 | 86.02 | 90.32 | 92.47 | 86.02 |

**Figure 1.**
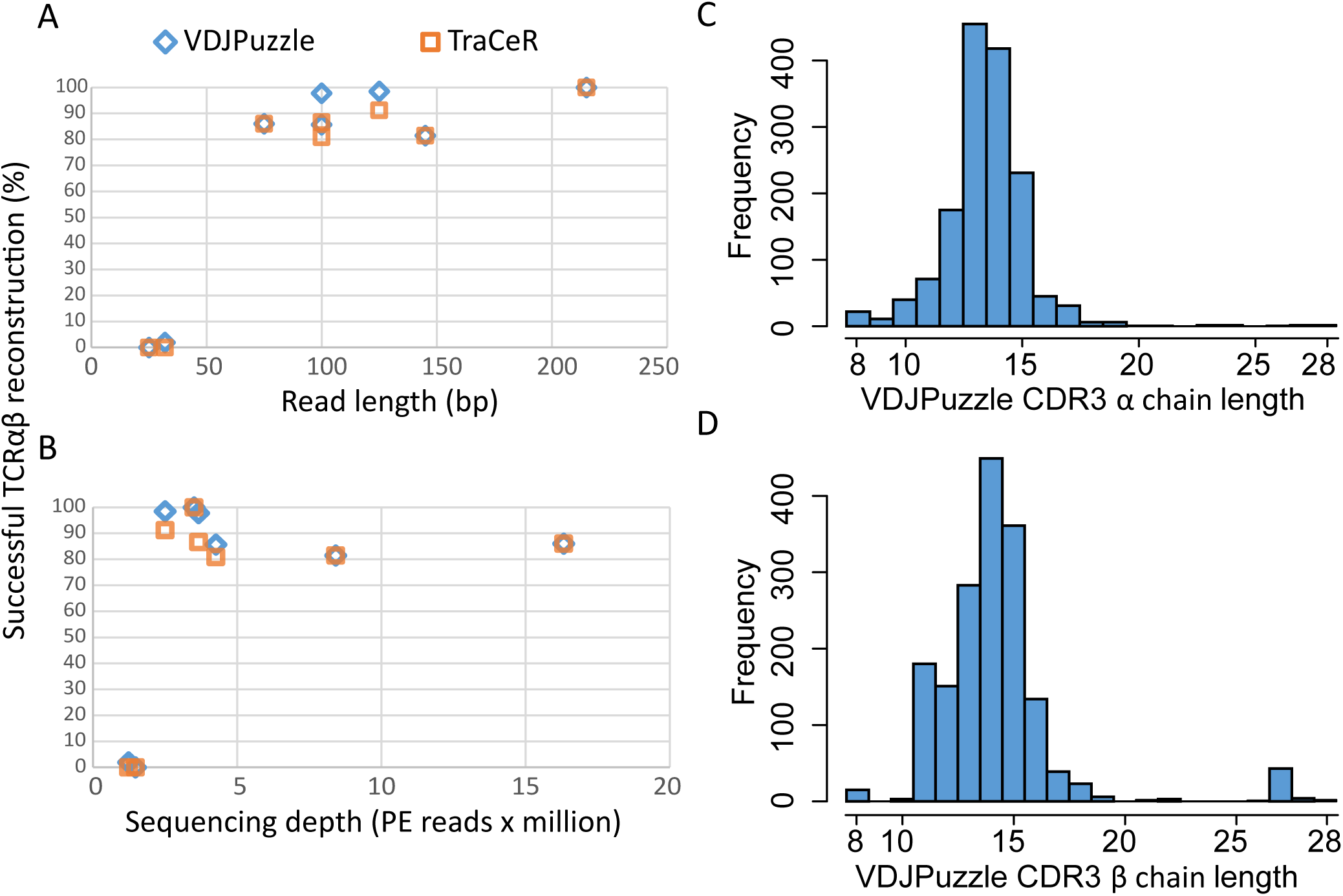
Success rates of TCRαβ reconstruction as a function of read length (A) and sequencing depth (B) using VDJPuzzle and TraCeR. Panels C and D show the distributions of the length of the reconstructed CDR3α and CDR3β regions from the VDJPuzzle output, respectively.

To further assess the quality of the reconstruction of TCRαβ sequence, we analysed the distribution of CDR3 amino acid sequences across both α and β chains, and the distribution of cells carrying double α chains. The average CDR3 length of the reconstructed TCRαβ sequences with VDJPuzzle was 14 amino acids for both α and β chains (Fig. 1C and D), with similar results using TraCeR (Fig. S1). This result shows a distribution of CDR3 lengths consistent with those previously estimated with other methods. In addition, we assessed the distribution of single cell carrying double α chains. One of the major advantages of using scRNA-seq to reconstruct TCR sequences is the possibility to detect double α chains within a single T cell. Overall, 30% (n=395) of the cells analysed here presented more than one α but not double β. In a single study (datasets 3 and 4 in Table 1), 43% (n=333) of the cells sequenced presented more than one α, and 44% (n=337) had more than one β sequence detected. Notably, 29% (n=225) of these cells had both more than two unique α and two unique β chain sequences, thus suggesting that in this study multiple cell could have been sorted in a single well. By filtering out cells with more than one α and one β, a total of 309 unique TCRαβ sequences were identified across all datasets. There was no clonotype (defined as cells bearing identical TCRαβ) overlapping between datasets.

### Effect of sequencing depth and read length on the TCRαβ detection using simulated datasets

To systematically investigate the effect of sequencing depth and read length, we generated simulated datasets with different sequencing depth and read length to assess the success rate of TCRαβ reconstruction. Simulated datasets were all derived from the original datasets 1 and 2, which had the deepest coverage (∼8.4 million PE reads per cell) and longest read length (Table 1). The original dataset consisted of a total of 54 single cells originated from HCV specific CD8+ T cells from a single subject that previously cleared HCV. Of these cells, 18 were directly sorted from peripheral blood mononuclear cells (PBMC-derived T cells) and the remaining 36 were sorted after *in vitro* expansion following stimulation with cognate antigen. Of these 36, 18 were sorted after a second antigen restimulation 24 hours prior to sorting ^17^). From each cell, we generated 16 randomly subsampled scRNA-seq datasets with all combinations of four different sequencing depths (0.05, 0.25, 0.625 and 1.25 million PE reads) and four different read lengths (25, 50, 100 and 150 bp) (Fig. 2A). For each of the 16 subsampled datasets, the TCRαβ sequence was reconstructed using VDJPuzzle ^17^, and the success rate was calculated (Fig. 2B and Fig. S2). Only TCRαβ sequences with a complete CDR3 recognised by the international ImMunoGeneTics information system (IMGT, ^25^) were considered as an exact TCRαβ reconstruction.

**Figure 2.**
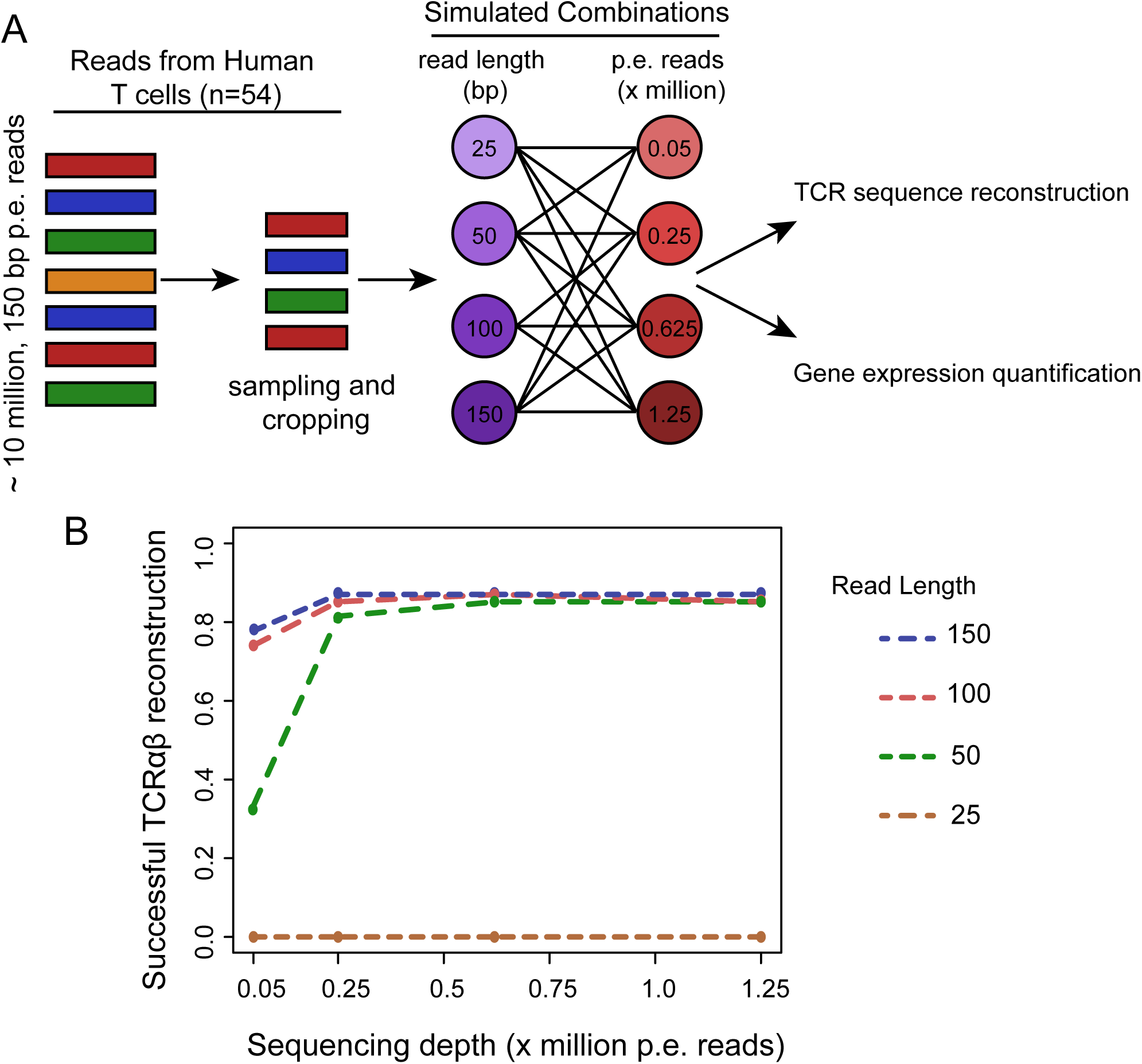
A) Generation of the simulated datasets from real scRNA-seq data 1. B) Success rate for TCRαβ reconstruction as a function of read length and sequencing depth from the simulated datasets.

Success rate of paired α and β was above 80% for datasets which had a minimum read length of 50 bp and a depth of at least 0.25 million reads. This rate was substantially diminished up to 0% for datasets with a number of PE reads per cell below 0.25 million PE reads (Fig 2B). Finally, the proportion of cells with double α detected was also proportional to both read length and sequencing depth, with the highest success rate corresponding to a depth of 0.25 million PE reads and a read length above 100 bp (Fig. S3). The relationship between the success rate of TCRαβ reconstruction and both sequencing depth and read length was fitted with a sigmoidal function (Fig. S2). The success rate in TCRαβ reconstruction from the experimental datasets (the real dataset) closely followed this specific relationship (*r*=0.97, *p* <0.01).

### The effect of read length and sequencing depth on the quantification of the gene expression profile

Next, we used the 16 subsampled scRNA-seq datasets to investigate the effect of sequencing depth and read length on read alignment and gene expression quantification. Surprisingly, we observed a slight increase in the total number of aligned PE reads in datasets with shorter read length, especially when the read length was below 100 bp (Fig. 3). This higher level of total read alignment at short read length can be attributed to an increased proportion of reads with multiple alignments, and more discordant alignment of PE reads (Fig. 3). Notably, this relationship with read length was also observed for the proportion of concordant pairs aligned, but with a lower proportion for reads of 25 bp long compared to 50 bp. We did not observe any effect of the sequencing depth on read alignment.

**Figure 3.**
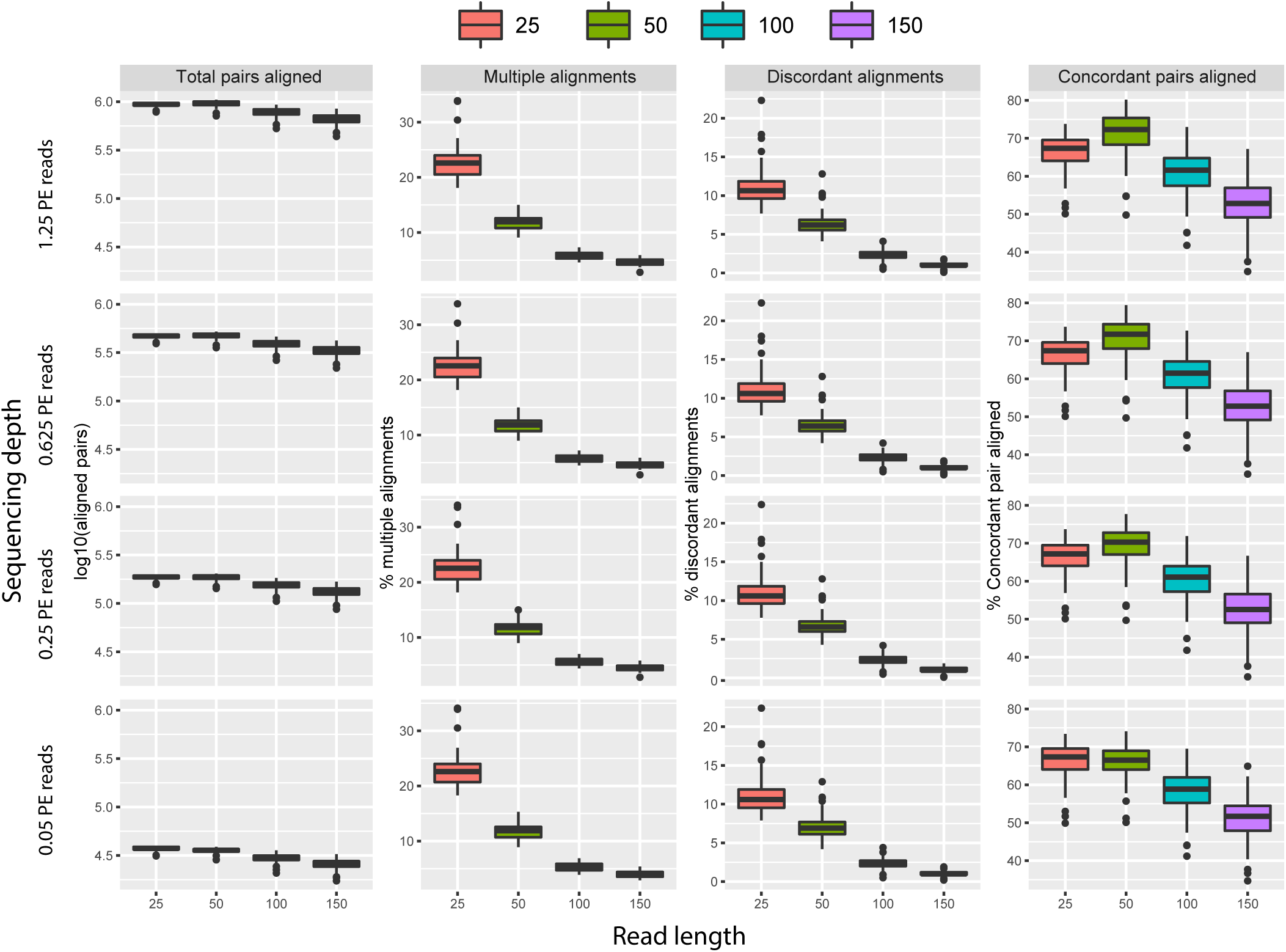
Analysis of the alignment of the simulated datasets as a function of sequencing depth and read length. Shown is the number of paired-end reads aligned (in log10 scale), along with the proportion of concordant and discordant pairs, and of multiple alignment instances.

To assess the effect of this trend on the quantification of genes, fragments per kilo base per million (FPKM) were calculated allowing only one alignment per read, hence eliminating a potential confounding factor of multiple alignments. We found that the number of detectable expressed genes (those with FPKM>1) was positively correlated with sequencing depth (Pearson correlation = 0.89) but negatively correlated with read length (Pearson correlation = -0.93). The number of genes that were expressed in at least 10% of the cells showed a similar correlation with sequencing depth and read length (Table 3, Fig. 4A). Notably, there was a positive relationship between number of genes expressed among cells within the same dataset and read length for sequencing depth smaller than 0.625 million PE reads, while there was no variation at higher sequencing depths (Fig. 4B).

**Table 3.** Number of genes expressed in at least 10% of the cells in the simulated data sets, comprised of subsamples of the scRNAseq data set 1, with various sequencing depths (columns) and read lengths (rows).

|  | Sequencing depth (PE reads x million) |  |  |  |
| --- | --- | --- | --- | --- |
| Read length (nt) | 0.05 | 0.25 | 0.625 | 1.25 |
| 25 | 6,081 | 8,497 | 9,184 | 1,0849 |
| 50 | 5,665 | 7,801 | 8,255 | 8,240 |
| 100 | 5,141 | 6,879 | 7,333 | 7,440 |
| 150 | 4,836 | 6,458 | 6,824 | 6,936 |

**Figure 4.**
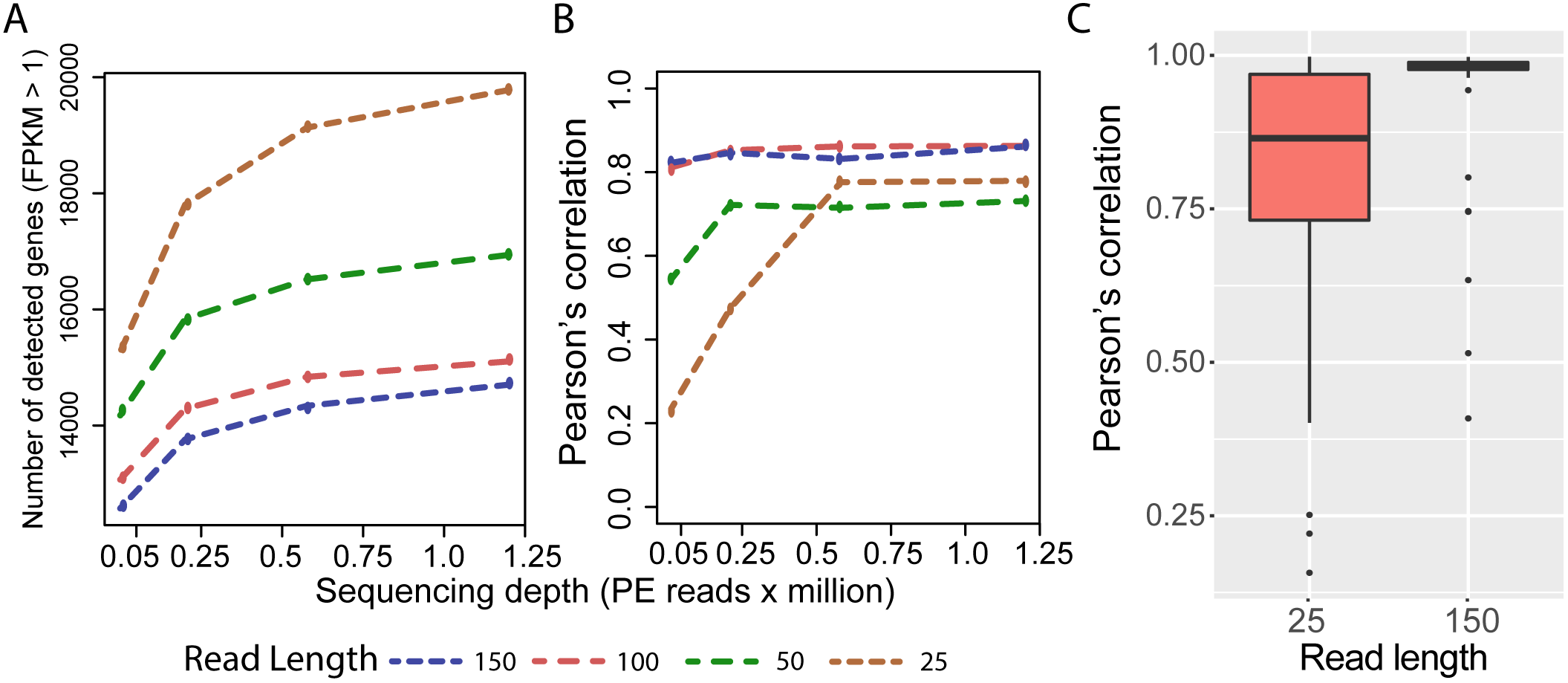
The effect of read length and sequencing depth on the technical error variability using simulated scRNA-seq datasets. A: Number of identified expressed genes (Fragment per Kilobase per Million reads; FPKM>1) as a function of read length and sequencing depth (A). B: Mean pairwise cell-to-cell Pearson correlation of gene expression values as a function of sequencing depth and read length. C: The distribution of pairwise cell-to-cell Pearson correlation of gene expression values using subsets of different read length drawn from the original dataset. Original dataset had a read length of 150 bp with depth >8 millions PE reads, two samples drawn from this dataset were taken, with length 25 bp and same depth.

In order to quantify the reliability of the gene expression profile as a function of read length and sequencing depth, two simulated datasets with a sequencing depth of 0.05 million PE reads were generated, with read length of 25 bp and 150 bp, respectively. Two replicates for each dataset were simulated. This analysis showed a significantly higher correlation between the gene expression profiles of paired cells from the two replicates with read length 150 bp when compared to the two replicates with read length 25 bp (Fig. 4C). This result suggested that gene expression profiles from short read length dataset have higher levels of technical noise.

To further assess how the technical variation generated by shorter read length and lower sequencing depth affects the identification of the three cell sub-populations available from the experimental scRNA-seq data of HCV specific T cells ^17^, a clustering algorithm was applied on all the simulated datasets. A newly developed bioinformatics tool CIDR ^26^ was used to perform dimensionality reduction, Principal Coordinates Analysis (PCoA) and clustering on the scRNA-seq gene expression profiles. When forced to identify three clusters (since there are three cell-types in the dataset), CIDR achieved the best clustering when the dataset has >= 100 bp long (Fig. 5A, B, Fig. S4). A higher misclassification rate with shorter read length was observed: 28% for read length 25 and 50 bp, and 9% for read length 100 and 150 bp (Fig 5C, Fig. S4). Sequencing depth did not affect the misclassification rate. To investigate whether the ‘tightness’ of the clustering is affected by sequencing depth and read length, the within-cluster-sum-of-squares of each cell type was computed. Consistent with the misclassification analysis, longer reads led to tighter clusters, reflected by a substantial decrease in within-class-sum-of-squares for PBMC derived Ag CD8+ T cells (Fig 5D, Fig. S4). The effect of read length was less pronounced for the other two in vitro expanded subpopulations, as these are biologically more close to each others when compared to the blood derived original population.

**Figure 5.**
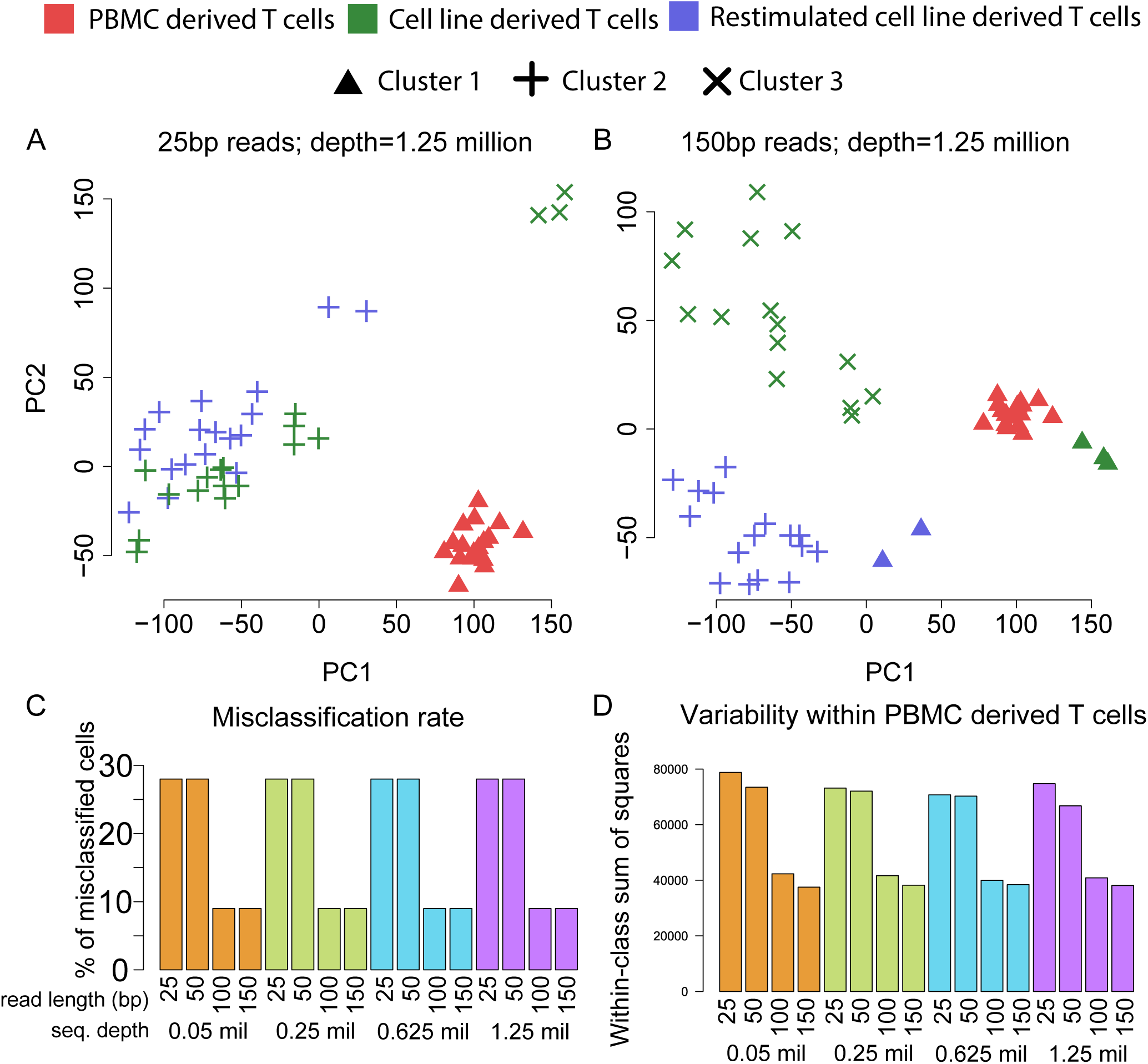
Clustering analysis for the three populations of HCV specific CD8+ T cells. Panels A and B display Principle Coordinate Analysis of the three subsets of cells by varying read length (25 to 150 bp). Coverage for each dataset was set to 1.25 millions of PE reads per cell. The point colours correspond to the ‘ground truth’ cell type labels (see legend), while the three point styles correspond to the three identified clusters (circle, triangle and cross). Clustering analysis was performed using CIDR, and forcing the number of clusters to be n=3. Panels C and D display the misclassification and the variability within the same cell type (within-class sum of squares) as a function of read length and sequencing depth, respectively. Panel D displays only results from PBMC-derived T cells.

To analyse the effect of read length and sequencing depth on specific gene categories, the distribution of gene expression levels (in terms of log(FPKM)) was analysed for highly expressed genes (average FPKM > 100), lowly expressed genes (average FPKM < 100), housekeeping genes, and transcription factors in all the subsampled simulated datasets. Independent of the gene category, there was a reduction in the number of genes identified with an expression level below 100 FPKM in datasets with a low sequencing depth (< 0.05 PE reads x million, Fig. S5). This effect was more evident among the transcription factors, where a combination or short read length and low depth led to a complete loss of lowly expressed genes. There was an increase in the frequency of highly abundant genes with the decrease of read length. To illustrate these trends, six individual genes were considered: two housekeeping genes (*GAPDH, RPL7A*, and *RPL34*), two genes constitutively expressed in CD8+ T cells (*CD8B* and *TRAC*), and one transcription factor (*GAS5*, which is associated with T cell proliferation ^27^). The analysis showed that, contrary to the expectation, the gene expression profile of the selected housekeeping genes varied significantly for low depth and short reads (Fig. 6). *GAPDH* and *CD8B* expressions were positively correlated with the read length, while a significant variability was detected for *GAS5,* independent of sequencing depth and read length. *TRAC* did not show any substantial variation.

**Figure 6.**
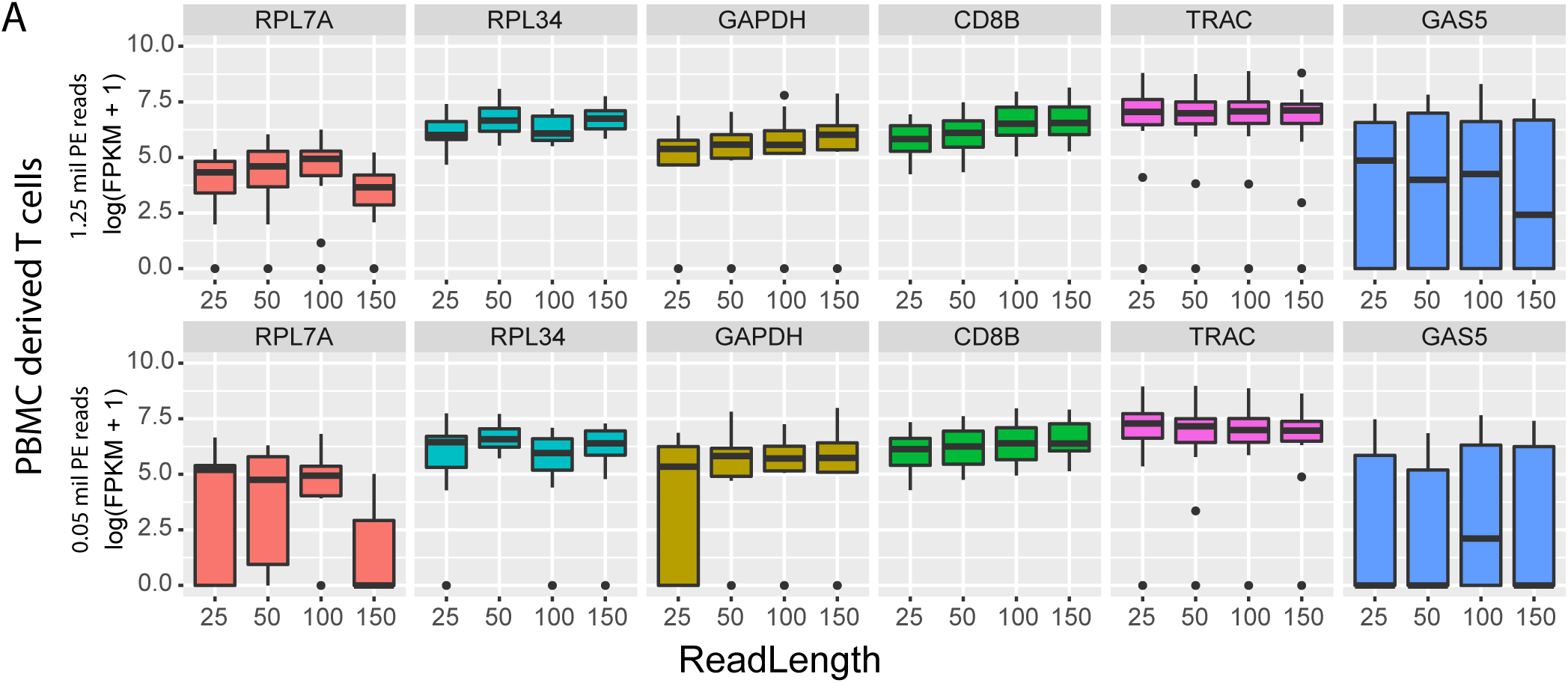
Gene expression profiles of selected genes identified from dataset 1, human HCV-specific CD8+ T cells (Table 1).

## DISCUSSION

This study aimed to explore how sequencing depth and read length of scRNA-seq dataset affect various downstream analyses, such as transcript reconstruction, gene expression estimation and cell-type identification. The overall messages of this study can be summarised with two major findings. Firstly, by combining available algorithms for TCRαβ detection, along with simulation-based analysis, this study has revealed that accurate detection of full-length TCRαβ is possible and achievable with sequencing depth below 250,000 PE reads, and with a minimum of read length of 50 bp. The detection rate of full length TCRαβ is at least 80% for reads with a sequencing depth >0.25 million PE reads of at least 50bp long. Secondly, the use of short reads (25 or 50 bp) is associated with a higher number of detected genes when compared to datasets with longer reads. However, this increase in gene expression quantification is also associated to a diminished accuracy and increased misclassification of cell populations. Hence, short read datasets are more prone to technical noise. The proposed analysis showed that despite accurate detection of full-length TCRαβ with short reads and relatively low sequencing depth, the gene expression profile is characterised by high level of technical variability. Future experimental designs should consider the quality of the reads as an important feature to obtain reliable results.

Analyses of simulated and real scRNA-seq data showed that current methods, such as SMART-seq2 are consistent with a capture efficiency between 3-10% of the total mRNA available ^8^. In the analysis proposed here, the effect of low sequencing depth in the quality of gene expression quantification and TCR reconstruction is likely to be associated to the poor library capture efficiency of mRNA from single cells (<10%) ^8^, hence it is conceivable that downstream analyses are not affected by large increase in sequencing depth. Low depth scRNA-seq has also been utilised to show that 50,000 reads are sufficient to classify a cell type in a sample of 301 cells ^12^. On the other hand, this may not be sufficient when more homogenous populations are involved, such as central memory and effector memory cells from the same antigen-specific repertoire. Deep sequencing of single cell library may still be required to improve detection of low abundant transcripts. Indeed, an important issue for scRNA-seq data is the very large amount of genes with zero expression ^6^. This observation results from real zero expression genes that a single cell may have at the time of RNA extraction, as well as dropout events, which are due to inefficient mRNA capture and library processing.

Full length TCRαβ can be accurately estimated and linked to the gene expression profile of the same cell. The analysis also showed multiple instances of single cells with at least two α and two β sequences detected. These findings are likely explained by the presence of multiple cells per well being sequenced, and the higher detection rate of double chains in the Th17 dataset (datasets 3 and 4) is likely due to the larger sample size compared to the other studies. The high success rate obtained with both available software programs further support the high quality of the scRNA-seq data, which significantly improve the quality of TCR reconstruction with more classical approaches such as bulk sequences and Sanger sequencing. Along with TCRαβ full-length data, the entire transcriptome can be interrogated to identify specific gene profiles associated to T cell subsets, along with the relationship with the TCRαβ clonotypes. Single cell approaches are therefore likely to increase further the accurate identification of novel markers, which could be utilised for detecting novel subpopulations of cells, for instance in flow cytometry. Another improvement is to introduce bar coding of the cell, with approaches such as MARS-seq ^28^. This novel approach however still lacks incorporation of full-length transcriptome sequencing, hence affecting the accurate detection of full length TCRαβ.

A drawback of current scRNA-seq approaches to study gene profiles is the relatively high cost ^3,4^. Currently, an Illumina HiSeq run can sequence up to 400 million PE reads of 150 bp in a single run. This allows generation of sufficient sequences reads for 640 single cells with depth of 0.625 PE reads with a total cost (library preparation + sequencing) of ∼US$10,000. This estimate assumes that the library preparation cost is approximately $5 per cell, which is achievable with in-house reagents. With these settings the expected number of reconstructed paired αβ TCR is about 85%. This cost is lower or equal to the cost of reconstructing TCRαβ from single cell using standard Sanger sequencing (without transcriptomic quantification). Barcoding approaches could significantly reduce the library preparation cost, down to <$1. However, this may impact the accuracy of TCRαβ detection as current bar coded methods have a 3’ bias with a significant loss of transcript depth on 5’ end reads ^3^. This gives us confidence that scRNA-seq technology can be scaled up to a large number of cells to comprehensively study the role of T cells in experimental models and human disease states.

## CONCLUSIONS

This study shed light on the effect of sequencing depth and read length of a scRNA-seq dataset on the quality and quantity of detection of gene expression profiles, and in particular of highly variable genes. High success rates in TCRαβ reconstruction are achieved with minimum depth and with reads >50 bp. In contrast, accurate detection of gene expression profiles and identification of cell subsets require longer reads to minimize technical noise. Future analyses should consider these effects to ensure reliable and accurate single cell transcriptomic profiling.

## MATERIAL AND METHODS

### ScRNA-seq data

Raw data were downloaded from NCBI’s Gene Expression Omnibus and ArrayExpress (Table 1).

### Generation of simulated datasets

Simulated data were obtained by generating subset of reads from Dataset 1 in Table 1 by randomly reducing read length and sequencing depth using an in-house python scripts. Original dataset consisted of 54 scRNA-seq data all with read length of 150 bp and sequencing depth of about 8.4 million PE reads. Sixteen combinations of read length and sequencing depth were considered: read length of 25, 50, 100 and 150 bp; and sequencing depth of 0.1, 0.5, 1.25 and 2.5 million. A new set of paired fastq file for each combination was then generated. For each dataset, reduced depth was obtained by randomly subsampling the original set of PE reads while the shorter read length was obtained by randomly cropping original PE reads.

### Gene expression quantification

PE reads were analysed for quality control using FastQC, and reads were trimmed using Trimmomatic ^29^.

Alignment of PE reads was performed with TopHat2. For the alignment, the default option was used, (https://ccb.jhu.edu/software/tophat/manual.shtml) which corresponded to allow for only one alignment per read. In case of multiple alignments of the same read, the primary alignment was considered.

Gene expression was estimated with the pipeline available in Cufflinks 2.2.1, utilising CuffQuant with parameter –max-frag-multihits equal to 1, which allows maximum one alignment per fragment. Gene expression quantification (in FPKM) was normalised with CuffNorm. Resulting FPKM values were loaded in R using the package Monocle ^30^.

Downstream analysis, which included Pearson’s correlation analysis, number of genes expressed, and gene expression analysis by gene categories, was performed with an in-house R script. Transcription factors and housekeeping genes have been selected from available list in the literature ^31,32^.

### Dropout rate and clustering analysis

Dropout analysis, principal coordinates analysis and clustering were performed using CIDR (Lin et al. 2017), which requires raw read counts as input data. The tool featureCounts was used to obtain the read counts, with the –*primary* option to allow only primary alignments. The *dropoutCandidates* Boolean matrix output by the *determinDropoutoutCandidates* method of CIDR is used to calculate the figures in Table 3 and Figure 5 – a gene is considered ‘expressed’ in a sample if the corresponding entry in the *dropoutCandidates* matrix has a value of *FALSE*.

For clustering, the CIDR parameters *nCluster* and *wThreshold* were set to be 3 and 6 respectively, while the other CIDR parameters were left as defaults. Within each cluster, the first two CIDR principal coordinates were used to calculate the distances between all pairs of samples, the squares of which sum to the within-class sum of squares.

Misclassification rate was used to evaluate the accuracy of clustering, which is defined as the number of misclassified cells divided by the total number of cells. To define misclassified cells, each CIDR cluster is associated with the ground truth cluster, which gives the biggest intersection, and those cells that are not in the intersection are counted as misclassified cells.

### Reconstruction of TCRαβ

TCRαβ of the downloaded dataset were reconstructed using VDJPuzzle ^17^ and TraCeR ^18^. The exact procedure was followed as previously reported ^17,18^.

The fit of the proportion of cells with successfull TCRαβ reconstruction as a function of read length and sequencing depth was performed using a two-dimensional sigmoid function implemented in the scipy package in python (the “curve_fit”)

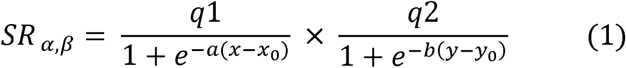

Where *x* represents the read length and *y* represents the sequencing depth. The obtained fitting values are *q*1 = 0.94, *q*2 = 0.95, *a* = 0.52, *b* = 98.67, *x*_0_ = 24.2, *y*_0_= 8.85.

## DECLARATIONS

### Ethics approval and consent to participate

Not applicable

### Consent for publication

Not applicable

### Competing interests

The authors declare that they have no competing interests.

### Availability of data and materials

The scRNA-seq data analysed are freely available and accession numbers are provided in Table 1.

### Authors’ contributions

SR analyzed and interpreted the scRNA-seq data. FL led the study and designed the experiments. SR, PL, JH, FL contributed softwares and performed bioinformatics analyses the data. SR, FL, PL and JH contributed to the statistical analysis, data collection, bioinformatics analyses. ARL, RB, AE contributed to the design of the study and in the analysis of the data. FL, SR, wrote the manuscript. VV JH AE were major contributors to the manuscript. All authors read and approved the final manuscript.

### Funding

We acknowledge the National Health and Medical Research Council of Australia (NHMRC) and Australian Centre for HIV and HCV Research (ACH2) for funding. SR is supported by the University International Postgraduate Award UNSW Australia. ARL is supported by an NHMRC Practitioner Fellowship (No. 1043067), AE is supported by an NHMRC Early Career Fellowship (No. 1130128). JWKH is supported by an NHMRC/National Heart Foundation Career Development Fellowship (No. 1105271), RAB, VV and FL are supported by an NHMRC Career Development Fellowship (No. 1060443, 1067590, 1128416).

